# The replisomes remain spatially proximal throughout the cell cycle in bacteria

**DOI:** 10.1101/097303

**Authors:** Sarah M. Mangiameli, Brian T. Veit, Houra Merrikh, Paul A. Wiggins

**Author notes:** To whom correspondence should be addressed. (P.A.W.) or (H.M.).

## Abstract

The positioning of the DNA replication machinery (replisome) has been the subject of several studies. Two conflicting models for replisome localization have been proposed: In the *Factory Model,* sister replisomes remain spatially colocalized as the replicating DNA is translocated through a stationary *replication factory.* In the *Track Model*, sister replisomes translocate independently along a stationary DNA track and the replisomes are spatially separated for the majority of the cell cycle. Here, we used time-lapse imaging to observe and quantify the position of fluorescently labeled processivity-clamp (DnaN) complexes throughout the cell cycle in two highly-divergent bacterial model organisms: *Bacillus subtilis* and *Escherichia coli*. Because DnaN is a core component of the replication machinery, its localization patterns should be an appropriate proxy for replisome positioning in general. We present automated statistical analysis of DnaN positioning in large populations, which is essential due to the high degree of cell-to-cell variation. We find that both bacteria show remarkably similar DnaN positioning, where any potential separation of the two replication forks remains below the diffraction limit throughout the majority of the replication cycle. Additionally, the localization pattern of several other core replisome components is consistent with that of DnaN. These data altogether indicate that the two replication forks remain spatially colocalized and mostly function in close proximity throughout the replication cycle.The conservation of the observed localization patterns in these highly divergent species suggests that the subcellular positioning of the replisome is a functionally critical feature of DNA replication.

**Author Summary:** Cell proliferation depends on efficient replication of the genome. Bacteria typically have a single origin of replication on a circular chromosome. After replication initiation, two replisomes assemble at the origin and each copy one of the two arms of the chromosome until they reach the terminus. There have been conflicting reports about the subcellular positioning and putative co-localization of the two replication forks during this process. It has remained controversial whether the two replisomes remain relatively close to each other with the DNA being pulled through, or separate as they translocate along the DNA like a track. Existing studies have relied heavily on snapshot images and these experiments cannot unambiguously distinguish between these two models: i.e. two resolvable forks versus two pairs of co-localized forks. The ability of replication to re-initiate before cell division in bacterial cells further complicates the interpretation of these types of imaging studies. In this paper, we use a combination of snapshot imaging, time-lapse imaging, and quantitative analysis to measure the fraction of time forks are co-localized during each cell cycle. We find that the forks are co-localized for the majority ( 80%) of the replication cycle in two highly-divergent model organisms: B. subtilis and E. coli. Our observations are consistent with proximal localization of the two forks, but also some transient separations of sister forks during replication. The conserved behavior of sub-cellular positioning of the replisomes in these two highly divergent species implies a potential functional relevance of this feature.

## Introduction

Rapid and faithful replication of the chromosome is essential for the proliferation of all living cells. Many bacteria possess a single circular chromosome. Replication is initiated at a single origin and two multicomponent protein complexes (replisomes) replicate bidirectionally around the chromosome, meeting at the terminus. Previous investigations of replisome localization have advocated two distinct models: In the *Factory Model,* the replication machineries at sister forks remain stationary and co-localized forming a *factory* through which the replicating DNA is pulled [1–6]. Alternatively, in the *Track Model*, the replication machineries translocate along a stationary DNA track, resulting in significant separations of the forks as replication progresses to sequences genomically distal from the origin of replication *(oriC)*, but re-localize as the forks converge at the terminus [7,8].

Although previous fluorescence microscopy studies have already reported on replisome localization, a significant weakness of these studies is that the statistical analysis relied heavily on still images (snap-shot data). As we will describe in this paper, without direct knowledge of the replication dynamics, it is difficult to differentiate between two pairs of co-localized replisomes that form as the result of re-initiation of replication and spatially-separated sister replisomes. In fact, both structures are observed in both *E. coli* and *B. subtilis.* Furthermore, the lengths of the replication and division cycles are highly variable in individual cells [9], creating the need for large-scale automated analysis which produces consistent results between time-lapse and snap-shot data.

Our method of analysis is based on the localization of the beta-clamp (DnaN) in both *E. coli* and *B. subtilis*. DnaN is a particularly informative proxy for the replisome complex because it is localized to the replisome [10] at sufficiently high copy-number that its position can be observed clearly throughout the cell cycle without significant photobleaching of the fluorescent label. Using time-lapse data, we tracked the progress of the replisome in individual cells over multiple cell cycles. Under slow growth conditions, the initiation and termination of replication can be observed explicitly by the assembly and disassemble of the DnaN foci.

Analysis of the time-lapse data reveals a significant confounding feature in the analysis of snap-shot images. For most of the cycle, sister replication forks maintain a sufficiently small separation such that the two foci cannot be resolved, but two sister forks are transiently resolvable in some cells, consistent with previous reports [11,12]. Irrespective of whether forks are co-localized or spatially resolved, they typically remain localized around midcell. This story is complicated by the phenomenon of re-initiation of replication before the end of the cell cycle [5]. Even under slow growth conditions, re-initiation predivision is observed in some cells. This uncoupling between the replication and division cycles leads to the appearance of DnaN foci localized to the quarter cell positions. Although this phenomenon is clearly observable in the time-lapse analysis, these foci can easily be misinterpreted as sister replication forks in snapshot analysis. However, careful statistical analysis of the snap-shot data clearly resolves two distinct subpopulations (resolved-sisters and colocolized replisome pairs). Our analysis of snapshot images produces results consistent with the replisome dynamics observed in time-lapse imaging, and can be applied to lower stoichometry replisome components, for which we find similar results. Therefore the analysis of both time-lapse and snap-shot images supports a model for replisome positioning in *E. coli* and *B. subtilis* where the sister replisomes localize with diffraction limited separation for the majority of the cell cycle.

## Results

### Time-lapse imaging of the replisome reveals proximal positioning

Replisome positioning was observed using time-lapse fluorescent microscopy by imaging fluorescent fusions to DnaN in both *E. coli* and *B. subtilis.* Cells were elongating exponentially with a doubling time of roughly 3 hours (Fig. 1), and multiple complete cell cycles were tracked under these relatively slow growth conditions.

**Fig. 1.**
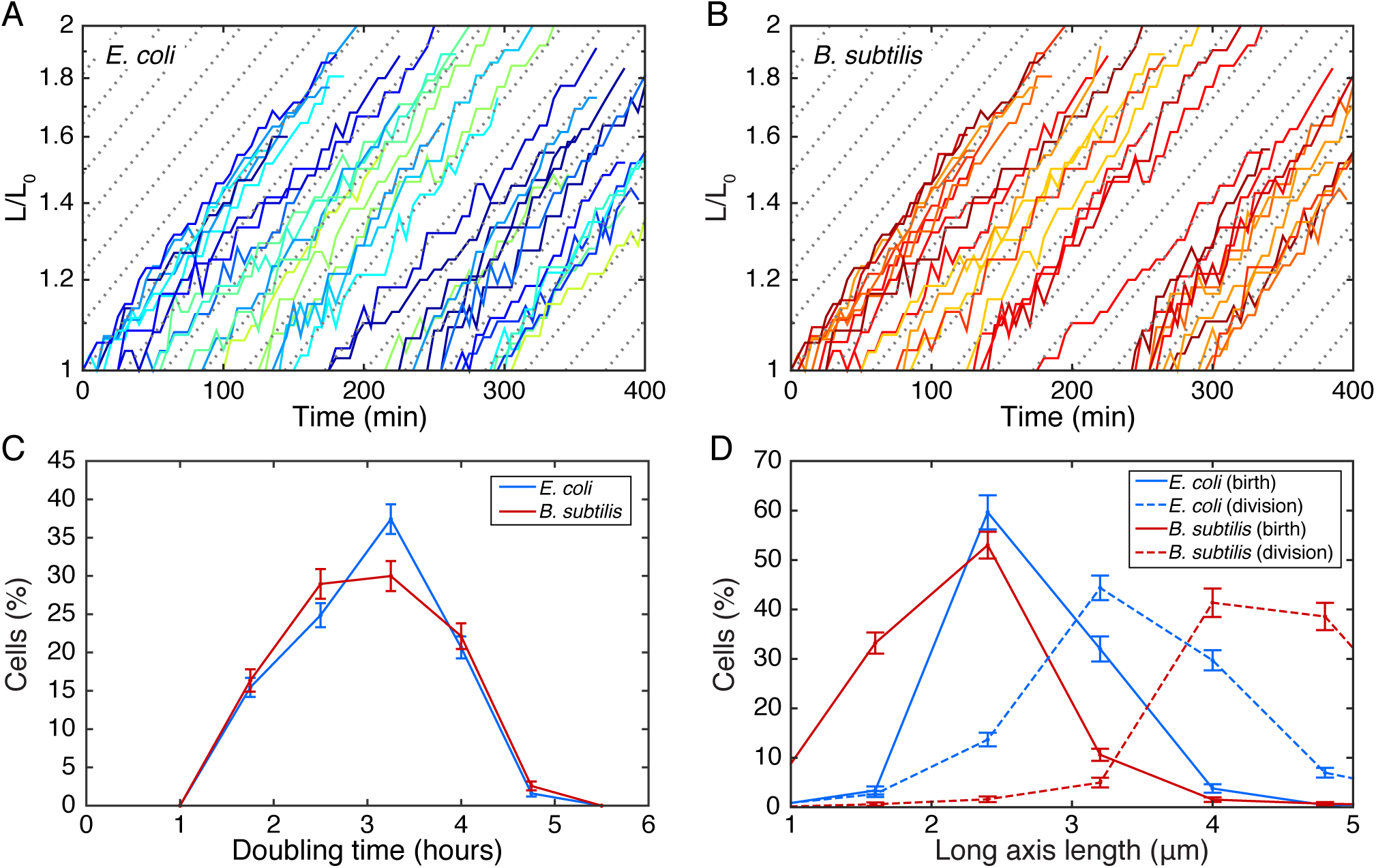
Characterizing cell elongation. (A-B) Typical length vs. time curves in *E. coli* and *B. subtilis*. Cell growth rate does not slow down significantly over time indicating that exposure to the laser is not damaging the cells. (C) Distribution of doubling times on microscope slide for *E. coli* and *B. subtilis*. (D) Distribution of cell lengths at birth (immediately following division of parent cell) and division.

It is widely accepted that the DnaN focus is localized to the replisome [10] and a good proxy for replication since: the beta-clamp is essential for replication, the focus is observed with midcell positioning (consistent with other replisome components), and the timing of assembly and disassembly is consistent with the known duration of replication.

A qualitative summary of the observed dynamics (see Fig. 2A) is as follows: In the absence of replication, there is no DnaN focus whereas actively replicating cells generally have between zero and four DnaN foci. DnaN foci tend to be localized either to midcell or the quarter cell positions, as has been reported previously [10,13] and is consistent with the localization of other replisome components. There is typically no focus or one focus at the midcell position. The typical temporal history of the position of a DnaN focus under slow growth conditions is concisely summarized by the kymograph in Panel B of Fig. 2: A focus appears. The position of the focus executes confined random motion. In some cells, the focus fissions into two dim foci which then fuse to re-form a single focus (of the initial intensity). The focus is then observed to disassemble. Typically there is a short period between the disappearance of the focus at midcell and either the roughly synchronous appearance of new foci at the quarter cell positions or cell division. DnaN foci were not observed to be intermittent: Once DnaN foci assembled they did not disappear and re-appear at the same cellular location.

**Fig. 2.**
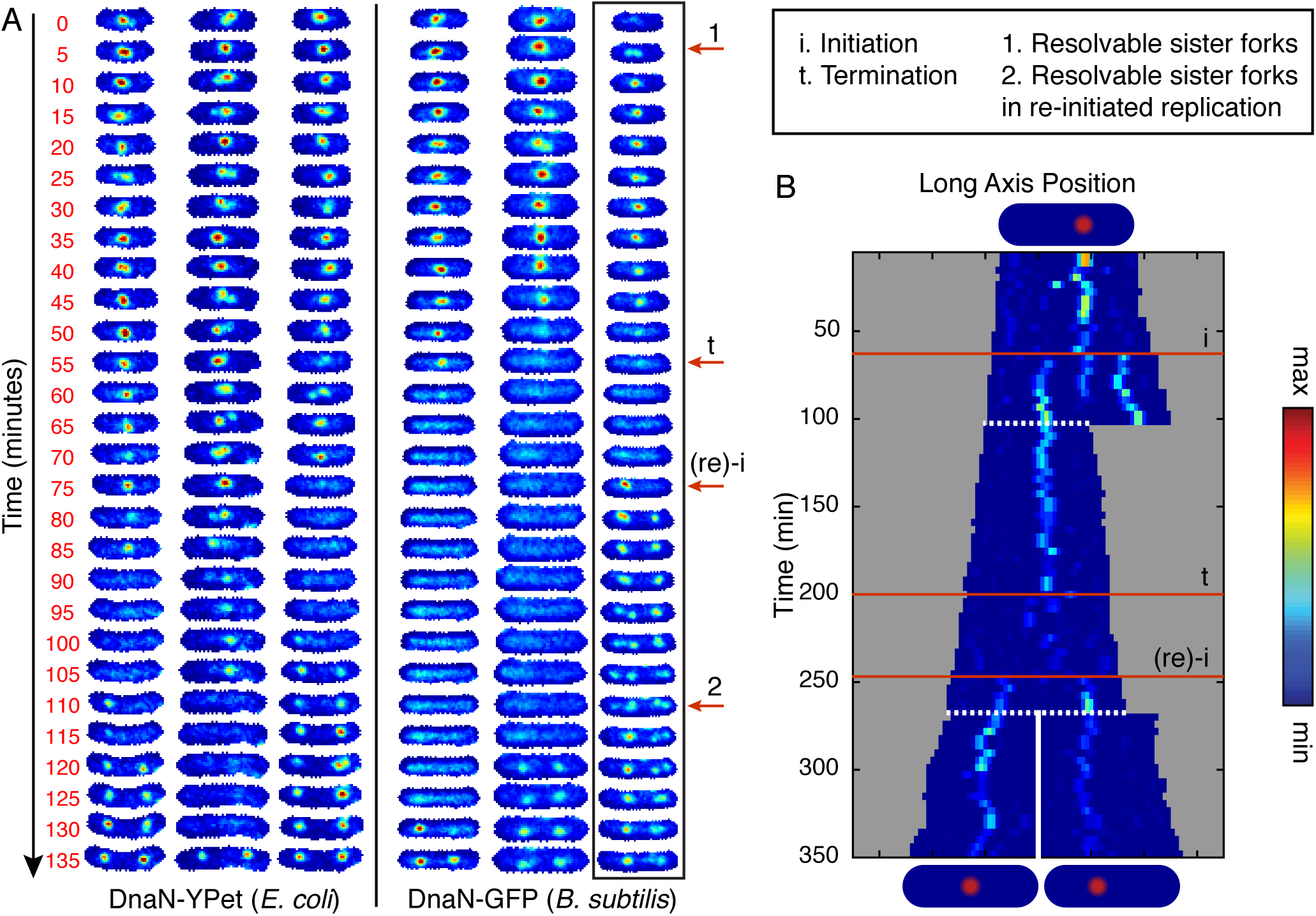
Visualizing replication forks in individual cells through complete cell cycles. A) Six representative examples of individual cells imaged at 5 minute intervals for DnaN-YPet in *E. coli* (left) DnaN-GFP in *B. subtilis* (right). Cells are tracked for complete cell cycles although images were cropped by up to a few frames to make the lengths consistent. Labeled red arrows point to example features in the boxed image strip. Starting at the beginning of the cell cycle, there is generally a single midcell focus representing both replication forks. However, occasionally sister forks can be resolved separately (e.g. arrow 1) but co-localize before termination of replication (e. g. arrow t). For a period of time, which varies cell to cell, no foci are observed until re-initiation on the newly replicated sister chromosomes (e.g. arrow (re)-i), an event which often happens before cell division. These new foci appearing at the quarter cell positions are consistent with replication factories since they can occasionally be resolved into sister replication forks (e.g. arrow 2). See also S1 Figure for additional full cell cycle images. B) Example single cell kymograph spanning multiple cell divisions for DnaN-YPet in *E. coli*. Cell images are projected along the long axis of the cell and stacked in sequence. White dashed lines indicate division events. Numbered red lines indicate initiation, termination, and re-initiation, respectively.

It is important to note the following qualifications about the number of DnaN foci: When the separation of foci are below the diffraction limit of our system (< 250 nm), only one focus will be resolved. Even when forks transiently separate enough to be resolvable, they remain well within the quarter cell positions: The average separation (when two foci are observed) is 0.2 cell lengths versus 0.45 cell lengths for foci near the quarter cell positions.

### Quarter cell foci are reinitiated replisome pairs

Single DnaN foci positioned at midcell are seen to fission and fuse (e.g. Fig. 2A, arrow 1). These midcell-positioned foci are always seen to disappear before cell division. On-the-other-hand, foci observed at the quarter cell positions can persist through a cell division, consistent with these foci representing re-initiation of replication. If instead these foci were separated pairs of sister forks, they would be expected to co-localize at the terminus and disassemble before the end of the cell cycle. Therefore qualitative analysis of the kymographs strongly supports a model where quarter-cell-localized foci each include pairs of reinitiated sister replisomes. In fact, these quarter cell foci are also seen to fission and fuse, occasionally allowing resolution of the individual sister replisomes (e.g. Fig. 2A, arrow 2).

To further test this model, we tracked DnaN foci in cells blocked for restart via a temperature sensitive version of the helicase loader protein, DnaC (*dnaC2* allele) [8]. Under the non-permissive conditions for the temperature sensitive mutant, the wild type cells were able to form quarter-cell-localized foci, however, the cells blocked for initiation were not (compare Fig. 3 panels A and C). To extend this analysis to many cells, we show conditional probability distributions of focus position given cell length in both the wild type and restart mutants. The absence of localizations near the quarter cell positions is clearly seen by comparison of Fig. 3, panels B and D. These data support our model that quarter cell foci represent reinitiated replication fork pairs.

**Fig. 3.**
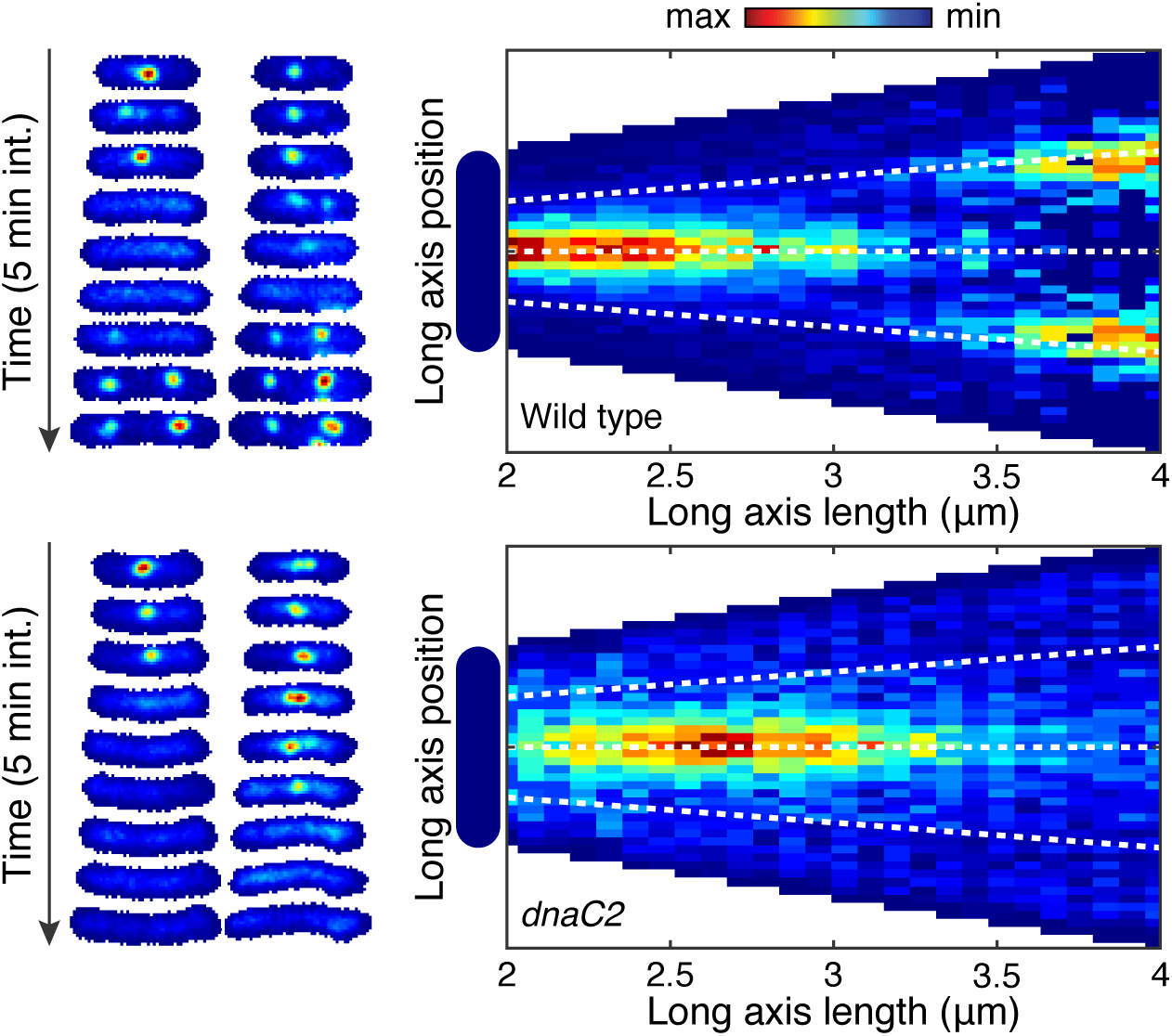
Blocked initiation leads to loss of quarter cell foci. DnaN-YPet is imaged in both in wild type and cells containing the *dnaC2* allele at 37°C. Under these conditions, cells containing the mutant allele will be blocked from initiating new rounds of replication. A) Example wild type cell towers showing the the disappearance of the midcell focus may be followed appearance of a pair of foci near the quarter cell positions. B) Conditional probability distribution (N = 4837 time points) shows localizations near the quarter cell positions in the wild type near the end of the cell cycle. C) Example cell towers for cells with blocked initiation do not show foci at the quarter cell positions after disappearance of the mid-cell focus. D) In cells blocked for initiation, conditional probability no longer shows a significant number of localization at the quarter cell positions (N = 1758 time points).

### Replication and division timing is asynchronous

In the event that re-initiation of the sister chromosomes happens before cell division (about 45% of the time under our conditions), we can only observe complete replication cycles if we analyze overlapping cell cycles. We visualize entire replication cycles using kymographs, where we project the cell images onto the long axis of the cell, and align the projections in sequence (See Fig. 2, Panel B). This representation confirms that for the majority of the replication cycle, the sister forks remain near mid-cell and usually cannot be resolved separately.

Since the timing of division is inferred from the analysis of the phase contrast image of the cell, some of the observed asynchrony could be accounted for by a failure to correctly segment the septum. However two lines of evidence refute this hypothesis: (i) Our previous work analyzing the cell-cycle dependent localization of FtsZ suggests that the timing of division is determined to a precision better than ±10% of the cell cycle in *E. coli* [14]. (ii) In this study, we never observed the midcell DnaN focus persist through cell division, consistent with accurate determination of cell division.

### Proximal replisome positioning is observed in snap-shot analysis

Due to significant cell-to-cell variation, it is essential to present statistical evidence for the classification of each focus as an individual replisome or colocalized replisome pair. We apply a fully automated analysis to characterize localization patterns for over 10,000 time points (detailed description included in the materials and methods section). For this analysis, foci are identified in the fluorescence image (S2 Figure) and precisely located within the cell. It is important to note that there was no hand selection of data. The dataset consists of all cells observed that were elongating and segmented without errors and therefore contains no investigator-based cell-selection bias.

We first investigate the focus positioning relative to cell length. In this analysis, cell length was used as a proxy for cell age because of *B. subtilis* chaining which makes detection of septum formation in phase images difficult. Cell length provided a consistent and reliable proxy for cell age in our analyses across both species and cell-length based analysis can be applied in snap-shot analysis where the cell age is unknown. The conditional probability density of focus position given cell length is shown in Fig. 4.

**Fig. 4.**
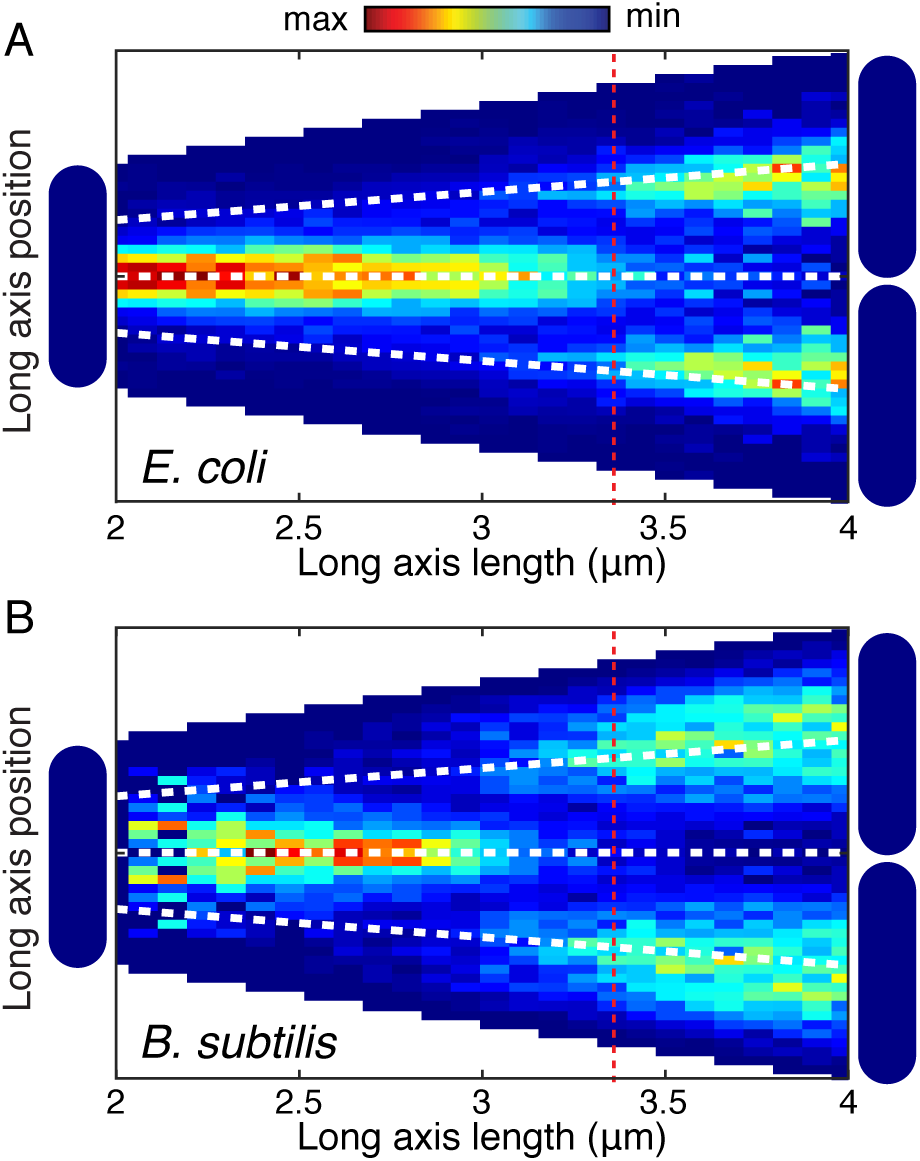
Focus localization pattern is similar in *E. coli* and *B. subtilis.* Probability of focus localization at a relative long axis position is shown as a function of cell length (analysis for over 10,000 foci in each organism). Dashed red line indicates mean length at re-initiation. A) DnaN-YPet in *E. coli* most probably localizes midcell until re-initiation when foci appear at the quarter cell positions. Dashed white lines indicate mid- and quarter-cell positions. B) DnaN-GFP in *B. subtilis* shows a similar localization pattern to DnaN-GFP in *E. coli* (Panel A).

The Factory Model predicts that foci are localized at midcell throughout the replication cycle and that re-initiation occurs at the quarter cell positions (if it occurs). In terms of the conditional probability, this model would predict a density blob at midcell which persists from early until late in the cell cycle. A second pair of blobs are expected to form at the quarter cell position corresponding to re-initiation in some cells. This second pair of blobs is expected to persist into the next cell cycle, each ending up in a new cell without the other. In contrast, in the Track Model, a blob is expected to begin the cell cycle at midcell, before splitting into two blobs (if the separation is high enough) before merging into a single blob again at midcell. The conditional probability data alone favors a factory-like model, but does not exclude the possibility of a Track Model, provided the separation of the forks remains very small as the replisomes translocate along the DNA.

Further analysis is informed by the typical cellular focus localization patterns shown in Fig. 5A. The relative frequencies of these localization patterns were quantified both overall, and by cell length. We find that short/young cells are generally observed to have a single focus. As the cells grow, the probability of observing two factories monotonically increases. In the Track Model, we expect the total number of two focus cells to grow significantly and then shrink as the forks first diverge from the origin and then reconverge at the terminus. This trend is not observed. Furthermore, before cells grow to a length where two foci is most probable, there is an increase in the probability of zero foci cells, consistent with the Factory Model where two foci cells have re-initiated but inconsistent with the Track Model where cells should transition from one to two foci without foci disassembling. Again, the Factory Model best summarizes the observed data.

**Fig. 5.**
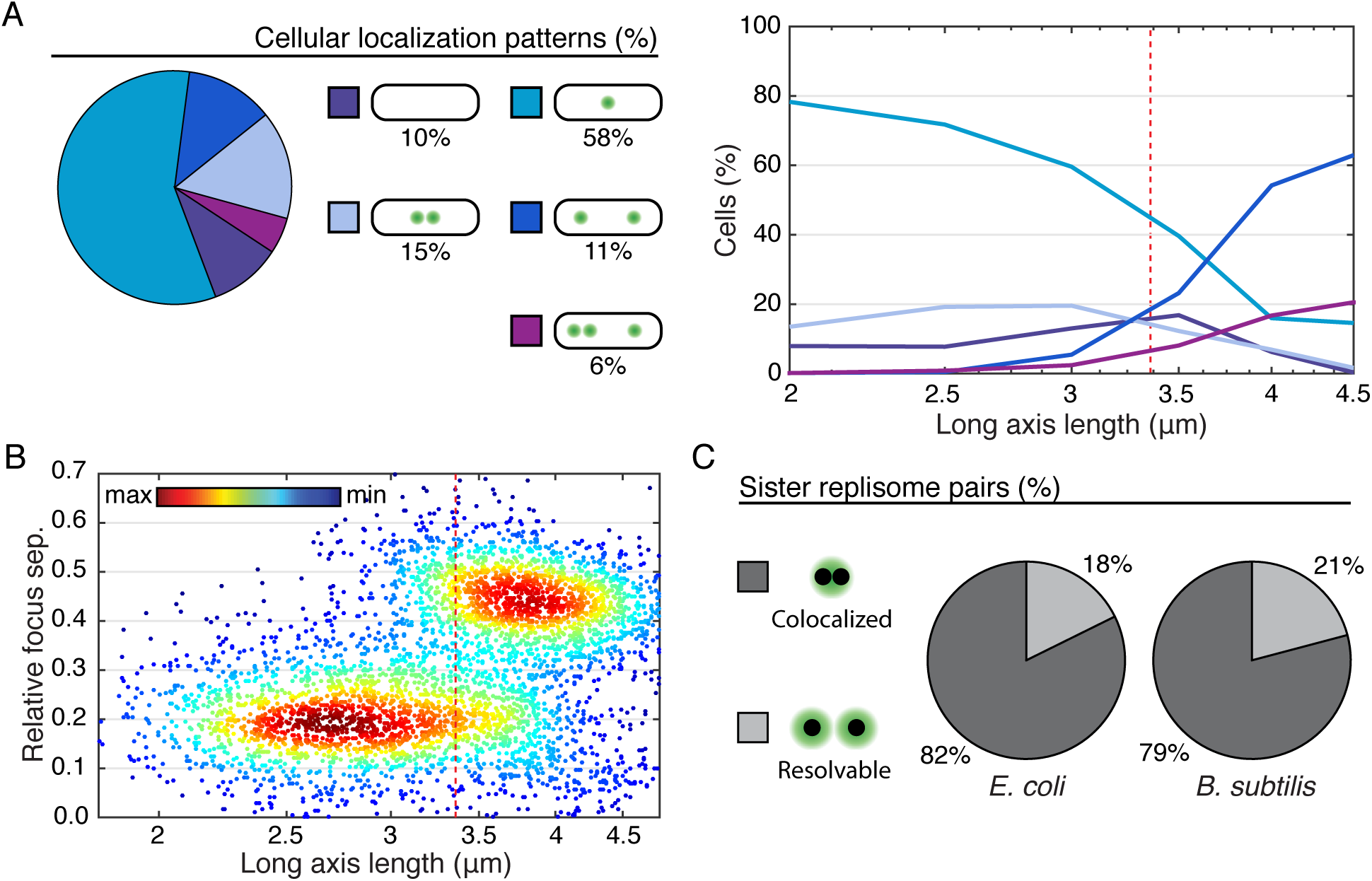
Characterizing cellular and subcellular localization patterns in large populations (> 10,000 time points). A) Left: Typical cellular localization patterns and their relative frequency of occurrence (in *E. coli*, see S4 Figure for corresponding *B. subtilis* data) are shown for all cells regardless of length. Extremely rare localization patterns (4+ foci) are not shown. Right: Relative frequency of the same typical cellular localization patterns as a function of cell length (age). Dashed red line represents mean cell length both at re-initiation and division. B) Multiple focus cellular localization patterns are distinguishable automatically based on inter-focus separations. Scatter plot shows relative focus separation as a function of cell length for multiple focus cells. Two well resolved populations are observed. Pairs of foci whose separation falls into the lower population are counted as individual members of a sister fork pair, whereas all other foci are each inferred to represent a colocalized replication fork pair. Dashed red line represents mean cell length at re-initiation and division. Example shown is for DnaN-YPet foci in *E. coli*. See also S4 Figure for corresponding *B. subtilis* data. C) Pie charts showing the fraction of sister replication fork pairs that are colocalized (replisomes have diffraction-limited separation) versus resolvable (each replisome appears as a member of a narrowly separated focus pair). In both organisms, replication fork pairs remain colocalized roughly 80% of the time, consistent with a factory-like model.

To automatically distinguish between the observed cellular localization patterns, it is essential to make a distinction between colocalized replication fork pairs and resolvable sister replication forks. Motivated by the qualitative observation that resolved sister forks rarely separate more than 0.2 cell lengths, we examine the joint probability distribution between focus separation (relative to cell length) and cell length shown in Fig. 5, Panel B. The joint distribution reveals that the population consists of two distinct sub-populations: In short/young cells, resolved foci are on average 0.2 cell-lengths in separation whereas in long/old cells, resolved foci are typically 0.45 cell-lengths in separation, consistent with quarter-cell positioning. The subpopulation model facilitates the identification of resolvable sister replication fork pairs. All pairs of foci whose separation is consistent with the lower separation population are counted as resolvable sister replication forks, whereas all other foci (including foci in single-focus cells) are inferred to represent a pair of colocalized replication forks (See S3 Figure and the associated methods section for more detail).

Considering cells of all lengths, we find that replication forks co-localize about 82% of the time for DnaN in *E. coli* (Fig. 5C). Statistical analysis was preformed similarly in *B. subtilis,* but because *B. subtilis* has a tendency to chain, division events are not always observable, at least not at the time of their occurrence. Since the division is not detected, more long cells that have re-initiated are observed. The exaggerated size of high separation population (S4 Figure) relative to the corresponding *E. coli* data is consistent with this known artifact. We also find that there is a longer time between termination of replication and re-initiation of replication on the sister chromosomes (D phase/G1 and G2 phase), increasing the fraction of zero focus cells. Strikingly, we find that the forks co-localize with 79% probability, roughly the same as observed in *E. coli* (Fig. 5C).

We applied the same statistical analysis used to quantify DnaN dynamics to snapshot images of three independent markers for the replisome (Fig. 6 and S5 Figure). Although DnaN is the only replisome component present at sufficiently high copy number to be imaged throughout entire cell cycles, the analysis we developed can be used to infer the dynamics of lower stoichiometry proteins based on snapshot images. We use SSB (single-stranded binding protein) and DnaQ (PolIII subunit) in *E. coli* and *DnaX* (clamp loader) in *B. subtilis* as additional markers for the replisome. Similar to that observed for DnaN, we again find that separated foci can be classified into two populations, one representing separated sister replisomes, and the other representing pairs of colocalized forks (S5 Figure, panel A). We count the number of foci in each cell (S5 Figure, panel B) and classify each focus as an individual replisome or replisome pair. It is important to note that the number of zero-focus cells is not particularly meaningful in this context as it depends on the growth rate of individual cells, and in particular, non-replicating cells cannot be excluded based on snapshot images. We find that sister replisomes colocalize with 74-85% probability (6), consistent with our observations based on DnaN.

**Fig. 6.**
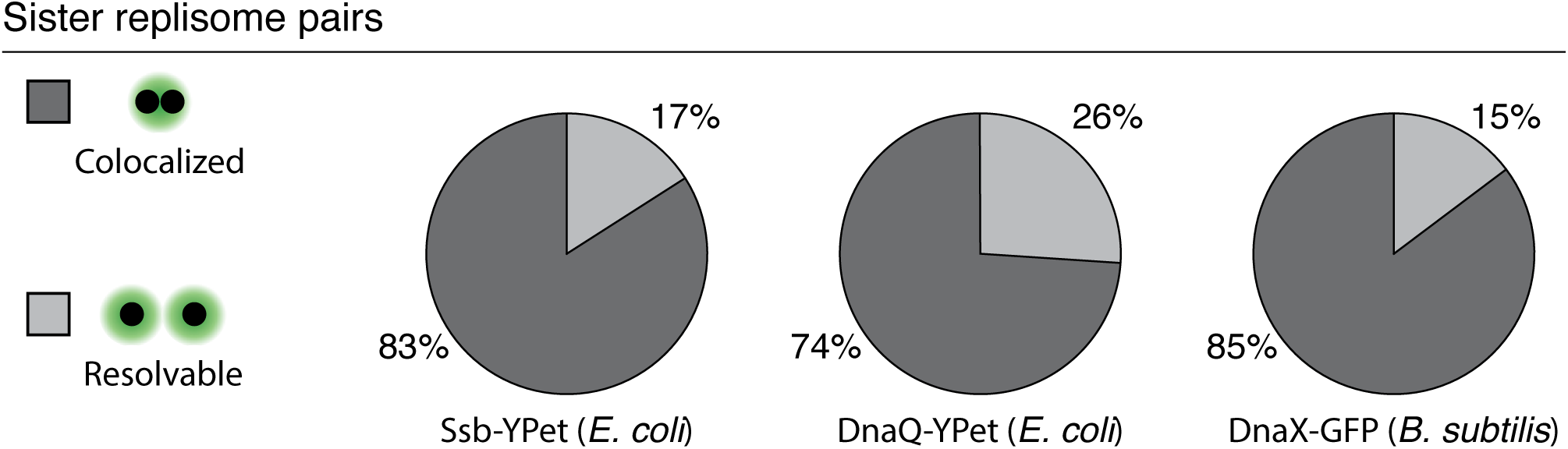
Additional markers for the replisome show consistency with DnaN. Pie charts showing the fraction of sister replication fork pairs that are colocalized (replisomes have diffraction-limited separation) versus resolvable (each replisome appears as a member of a narrowly separated focus pair) in SSB-YPet (N = 6493 Cells), DnaQ-YPet (N = 2187 cells), and DnaX-GFP (N = 10573 cells). Results are consistent with DnaN time lapse imaging. See also S5 Figure for complete analysis.

**Fig. 7.**
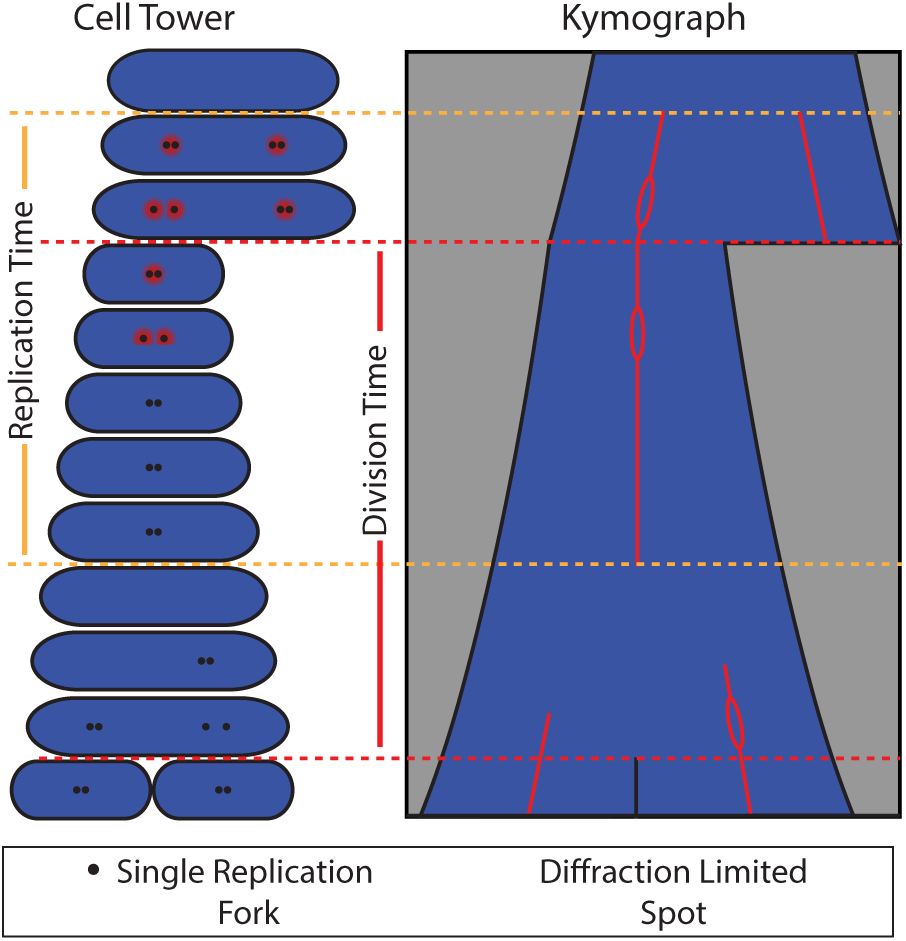
A schematic model for replication fork localization during complete cell cycles. Both the kymograph and cell tower representations are shown. Our model for replication fork localization is as follows: (i) The replication and division cycles are generally out of sync, so that full replication cycles occur across multiple division cycles. (ii) During replication, forks co-localize to a diffraction limited spot about 80% of the time, consistent with factory-like behavior. (iii) When the forks are individually resolvable, separation is small (< 0.33 cell lengths) and transient. (iv) Termination of replication is indicated by disappearance of foci. (v) Re-initiation often occurs before cell division near the quarter cell positions. Because these foci occasionally split into pairs of resolvable foci, they are consistent with factories containing two replication forks.

## Discussion

### Sister replisomes have proximal positioning

Both time-lapse and snapshot analysis strongly support a model where the two replication forks remain proximal throughout the cell cycle. At the beginning of the replication cycle, we always observe a single focus. We associate this event with co-localization of the forks at or near *oriC*. After initiation, the replisomes remain well within the quarter cell positions, usually co-localizing, but separating enough to be individually resolvable approximately 20% of the time. The observed midcell positioning of the replisome is consistent with previous reports from our lab suggesting that chromosomal loci in *E. coli* are localized near midcell immediately prior to duplicating [6]. Together this study and the aforementioned previous reports suggest that movement of the DNA through a relatively stationary replisome.

The occasional separation of sister replisomes suggests that, while the Factory Model correctly predicts the cellular-scale positioning of forks in both *E. coli* and *B. subtilis*, some elements traditionally associated with the Track Model are also correct in the sense that the sister forks need not be continuously colocalized. However, this must be qualified by noting that roughly 80% of sister replisome do appear to be colocalized by conventional fluorescence microscopy, and it is unlikely that this would be the case if the replisome translocating along the DNA were the complete model. This quantitative picture will be essential in interpreting short-time scale experiments which do not capture the cell cycle dynamics. Under our conditions, at least 80% of observed foci are in fact pairs rather than individual replisomes.

Our data strongly support a factory-like model, but do not address whether there is indeed a “factory”. In the strictest definition of a Factory, sister replisomes are coupled by a physical linker. While we do not exclude the possibility of a physical linker, such a linker would need to allow the replisomes to occasionally uncouple, or be sufficiently long to account for the observed separation events. The physical mechanism by which sister replisomes remain spatially proximal is an intriguing problem for future studies.

### Two types of focus separation and dynamics

We see two distinct types of separation and dynamics. In the first class of separation, the relative distance between the foci is under a third of the cell length, and is about a fifth of the cell length on average. Foci with these small separations always merge. In contrast, if re-initiation precedes division, we observe half cell length focus separation. In addition to having a larger separation, these foci are never observed to merge and ultimately end up in different cells. Furthermore, widely separated foci are unable to form in conditional mutants where replication restart is prevented. This localization behavior indicates that the half cell-length foci we observe pre-division are new pairs rather than individual replisomes.

### Replisome positioning dynamics are conserved in Gram-positive and Gram-negative bacteria

The comparison of replisome dynamics between E. *coli* and *B. subtilis* was particularly important given that the Factory Model was formulated based largely on data obtained from *B. subtilis* studies whereas the Track Model was formulated based predominantly on data from E. *coli* [7, 8]. Because of this, the reason for the existence of conflicting models was often attributed to the differences between the two species. Our data suggest that the conflicting models likely originated from differences in analysis and interpretation of data rather that species-specific replisome dynamics.

In every stage of the analysis, the replisome positioning data reveals striking similarities between *E. coli* and *B. subtilis*. Gram-negative and Gram-positive bacteria are highly divergent. Although the basic principles of replication are conserved, many differences exist between E. *coli* and *B. subtilis* replication, including regulation, the leading and lagging stand polymerases, and the loading of the replicative helicase. The data presented here reveal that replisome positioning is one of the conserved features, suggesting that there may be fundamental mechanistic reasons for precise subcellular localization of this complex. Future studies should reveal the mechanism by which the two forks remain proximal throughout the replication cycle and identify the underlying reasons (if any) for this conserved localization pattern.

## Materials and Methods

### Strain construction and growth

See Table 1 for strain list. PAW1181 was produced by P1 transduction of the mutation from PAW542 into PAW914. Cells were cultured overnight at 30°C in minimal medium with shaking. Prior to imaging, cells were set back to OD_600_ 0.1 and allowed to grow to OD_600_ 0.3. To ensure sufficiently slow growth that initiation would happen only once per-division cycle, we use minimal medium supplemented with only the essential nutrients. For *E. coli*, cells were cultured in M9-minimal medium (1X M9 salts, 2 mM MgS0_4_, 0.1 mM CaCl_2_,0.2% Glycerol, 100 *μ*g/ml each Arginine, Histidine, Leucine, Threonine and Proline and 10 *μ*g/ml thiamine hydrochloride). *B. subtilis* was cultured in Minimal Arabinose Medium (1x Spitzizen′s salts (3 mM (NH_4_)_2_S0_4_, 17 mM K_2_HPO_4_, 8 mM KH_2_P0_4_,1.2 mM Na_3_C_6_H_5_0_7_, 0.16 mM MgSO_4_-(7H_2_O), pH 7.0), 1x metals (2 mM MgCl_2_, 0.7 mM CaCl_2_, .05 mM MnCl_2_,1 *μ*M ZnCl_2_, 5 *μ*M FeCl_2_,1 *μ*g/ml thymine-HCl), 1% arabinose, 0.1% glutamic acid, 0.04 mg/ml phenylalanine, 0.04 mg/ml tryptophan, and as needed 0.12 mg/ml tryptophan).

**Table 1.** Strain List

| Stran No. | Genotype | Species | Reference |
| --- | --- | --- | --- |
| PAW914 | YPet-DnaN | *E. coli* | [15] |
| PAW885 | dnaN-gfp | *B. subtilis* | [10] |
| PAW542 | dnaC2 thrA::Tn10 | *E. coli* | [8] |
| PAW1181 | dnaC2 YPet-DnaN | *E. coli* | This study |
| PAW843 | ssb-ypet | *E. coli* | [15] |
| PAW913 | dnaQ-ypet | *E. coli* | [15] |
| PAW884 | dnaX-gfp | *B. subtilis* | [10] |

### Microscopy slide preparation

For imaging, we gently heat the appropriate growth medium with 2% by-weight low melt agarose (Fisher: 16520050). The agarose mixture is then molded into a thin rectangular strip (2mm X 4mm X 0.05 mm) and allowed to dry. One micro-liter of OD_600_ 0.3 liquid culture is spotted centrally on the pad. Once the spot has dried, a cover glass is placed over the pad and sealed around the edges with VaLP (1:1:1 Vaseline, Lanolin, and paraffin mixture). This leaves a small channel of air around the pad, particularly important for the growth of *B. subtilis*.

### Microscope Configuration

Imaging was performed on our lab-built inverted fluorescence microscope. Cells were imaged through a Nikon CFI Plan Apo VC 100x 1.4 NA objective. A retractable external phase plate (Ti-C CLWD Ph3 Annulus Module) was inserted into the light path during phase-contrast imaging but removed for fluorescence imaging to avoiding decreased signal due to the neutral density annulus on the phase plate.

For fluorescence imaging, we excite GFP and YPet proteins using a Coherent Sapphire 50 mW 488 nm or 150 mW 514 nm CW laser (see table 2). The beam diameter is expanded, providing uniform illumination over the field of view. An Acousto-Optic Tunable Filter (AOTF, AA Opto-Electronic AOTFnC-400.650) controls the laser excitation intensity. Images were collected on an iXon Ultra 897 512x512 pixel EMCCD camera. The microscope system is controlled by Micro-Manager.

**Table 2.** Microscope settings

| Protein | Laser Power (mW) | Exposure (ms) |
| --- | --- | --- |
| GFP | 1.1 | 600 |
| YPet | 0.93 | 600 |

### Imaging

Cells are imaged at about 25°C in both phase contrast (to determine cell boundaries) and fluorescence (to measure replisome position) at five minute intervals. The built-in software autofocus is used to ensure focus at each time point. Laser intensity is maintained as low as possible to avoid bleaching the fluorescent protein and damaging the cells. We are generally able to image cells for about four hours before photobleaching becomes too significant to reliably track the forks. For lower stoichoimetry replisome components, snapshot imaging was used where one phase contrast and one fluorescence image was taken at each field of view.

Non-permissive conditions for the temperature sensitive initiation mutant were achieved using an objective heater (Bioptechs, 38°C set point). Cells were placed on the heated objective about 10 minutes prior to the start of imaging. Imaging continued with frames at 5 minute intervals for several hours, long enough to capture the beginnings of subsequent replication cycles in wild-type cells.

### Characterizing cell elongation during microscopy

Using the cell boundaries determined from segmentation of the phase image, we track the long-axis length over time. Fig. 1A and B show typical length vs. time curves for *E. coli* and *B. subtilis*. Importantly, the cells were elongating exponentially (curves appear linear on a log scale) and the growth rate (slope) appears constant over time, indicating cell growth was not affected by prolonged exposure to the laser. We determine the doubling time for each cell by least squares fitting of the length vs. time curve with an exponential of the form:

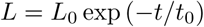

Where the parameter *t*_0_ was allowed to vary and *L*_0_ represents the length of the cell immediately following division. The doubling time was taken as:

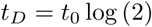

The distribution of cell doubling times for individual cells is shown Fig. 1C.

### Image Processing

Cells were imaged in both phase-contrast and fluorescence at five minute intervals. Higher time resolution led to significant photobleaching and/or slowed cell elongation. Imaging continued sufficiently long to capture at least one full cell cycle, about 3.5-4 hours. For snapshot imaging, only one phase-contrast and fluorescence image were taken at each field of view. The process used to analyze the phase and fluorescence images is outlined below.

### Cell segmentation from phase images

At each time point, we capture a phase-contrast image for the purpose of segmentation, the process of analytically identifying cell boundaries. Using our lab’s previously described custom segmentation tool (avalible at http://mtshasta.phys.washington.edu/website/ssodownload.php) [16,17], *superSegger*, we generate cell masks in each frame for further analysis and track cells frame-to-frame for entire cell cycles. In order to ensure complete cell cycles in individual cells, we must observe two division events. Cells are tracked starting from their birth (division of the parent cell) until division. The lengths of cells at birth and division are highly variable, and the distributions are shown in Fig. 1. We note that the distributions for *B. subtilis* is artificially shifted towards longer lengths because chaining often prevents visual identification of a septum until significantly later than the cell division event.

### Locating foci in fluorescence images

To manage xy drift, each phase image was aligned against the previous frame. The the corrected alignment of each phase image (and corresponding fluorescence image) was retained for further analysis. Foci are then identified in each fluorescence image by the process described below. A one pixel radius Gaussian blur is applied to each frame, after which we apply a watershed process to the conjugate image to identify regions around each intensity maximum. Regions external to the cell masks are excluded from further analysis. Regions internal to the cell masks will be called intensity regions. In order to precisely locate foci within each intensity region, we form a 3 pixel radius circular region at the location of the maximum-intensity pixel. Within the circular region, we model the intensity profile as Gaussian distribution with the following form:

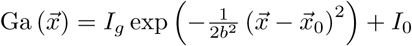

Here, the parameter *I_g_* defines the Gaussian peak amplitude and *I*_0_ represents the background intensity. The calculated focus position is 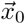 and the parameter *b* characterizes the focus width.

The focus position obtained from the Gaussian fit is used for all localization calculations. In addition, quantifying focus intensity is important for avoiding false detection events. As a measure of focus intensity, we sum the intensity in an 3 pixel radius disk centered at the focus location obtained from the Gaussian fit. The background intensity (*I_b_*) is defined as the minimum pixel value within the circular disk multiplied by the area of the disk. The standard deviation of intensity (*δI* respectively) is taken over the entire cell mask.

### Focus scoring

The statistical significance of a focus is measured by its score *σ*. The integrated background subtracted intensity (*I_a_*) inside a three pixel radius region is computed. The score is defined as:

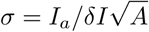

Where A is the area of the circular region over which the intensity was integrated. The factor 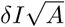 is the expected standard deviation in the integrated intensity over the integration area, assuming the noise at each pixel is uncorrelated. Foci scoring 4 or higher were retained. Lower scoring foci localized randomly throughout the cell and were consistent with stochastic fluctuation in the intensity (S2 Figure).

### Data Selection

Data processing was completely automated, with no selection by the investigator. Cells were selected for analysis based on the following three criteria:

- No segmentation errors
- Cells were growing exponentially with a doubling time between 1.5 and 4.5 hours.
- The cell length at birth was between 1-4 *μ*m.

The second criterion did not apply to snapshot images where the doubling time could not be calculated, and the last criterion was included mainly to prevent the analysis of long chains of *B. subtilis* cells. Additionally, each experiment was repeated on at least two separate occasions, and data was pooled for multiple experiments.

### Statistical analysis of focus localization

#### Calculation of focus separation

All possible unique distances between the foci in each cell were calculated, and the long axis components were retained. Relative separation is the long axis component of the separation between foci divided by the long axis length of the cell. Visualization of relative focus separation as a function of cell length resulted in two roughly Gaussian populations. We interpret the low separation population (*P_L_*) with transiently separated sister replication forks. The higher separation population (*P_H_*) is less straight-forward to interpret since these separations may be between pairs of re-initiated factories, a re-initiated factory and a member of a resolvable fork pair, or members of resolvable fork pairs on opposite sides of the cell. Additionally, this higher separation population is exaggerated in *B. subtilis* since chaining often prevents visualization of the septum. We fit a two-Gaussian mixture model to the observed distribution of separations using a maximum likelihood process where the Gaussian means, variances, and mixing fractions were allowed to vary.

#### Calculating probability of a replication factory

All cells with a single focus are interpreted to have a colocalized replisome pair. For multiple-focus cells, we identify resolvable sister replisomes by identifying pairs of foci whose separation is most probably associated with *P_L_* (as determined by the posterior probability distribution resulting from the Gaussian mixture model). For each cell, we then define the number of colocalized replisome pairs as the total number of foci minus twice the number of resolvable replisomes. This process is illustrated in S3 Figure.

## ACKNOWLEDGMENTS

This research was supported by NSF grant MCB1243492. The funders had no role in study design, data collection and analysis, decision to publish, or preparation of the manuscript.

## Supplementary figures

**S1 Figure.**
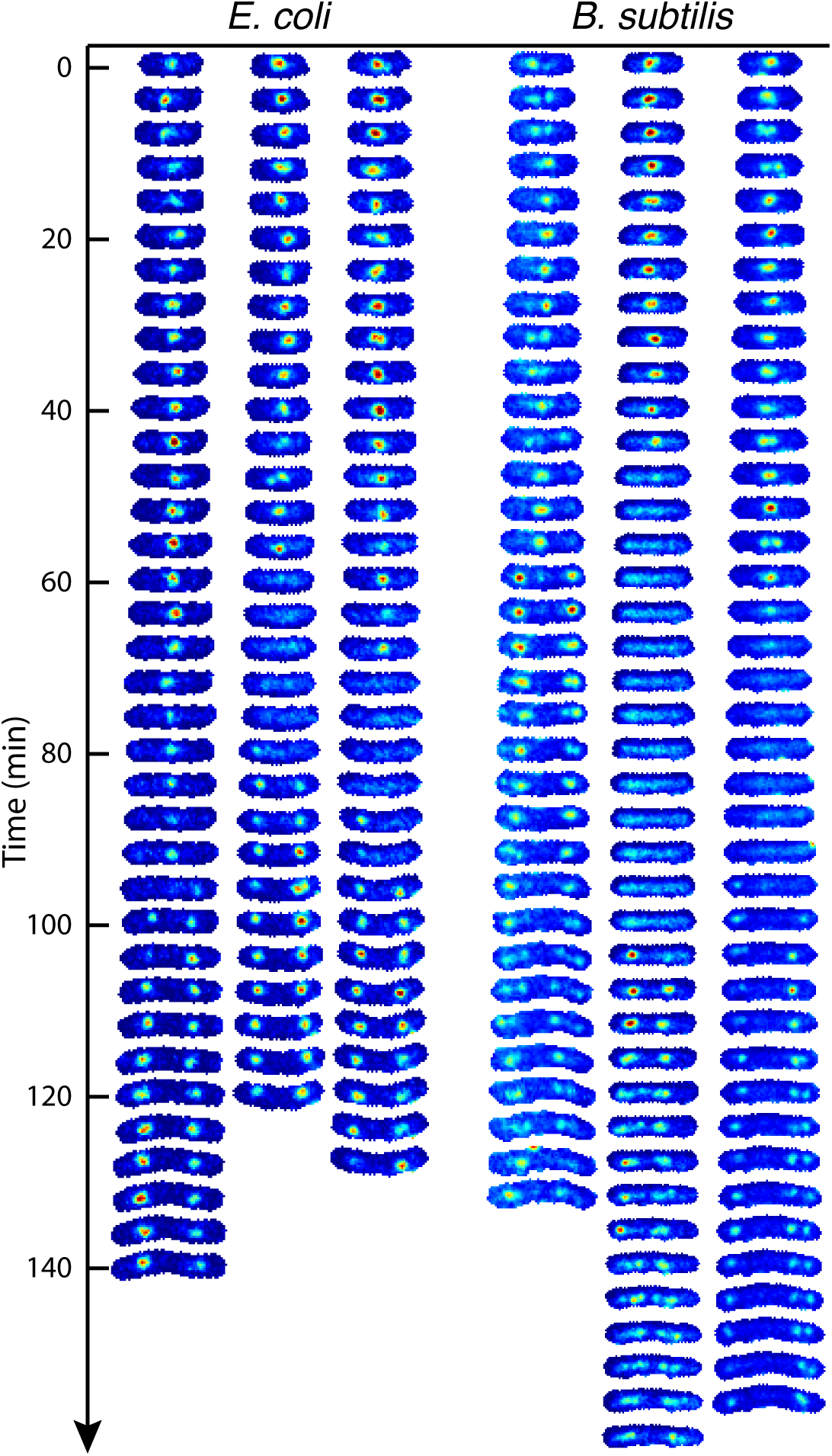
Additional cell towers. Additional example cell towers for both *B. subtilis* and *E. coli*.

**S2 Figure.**
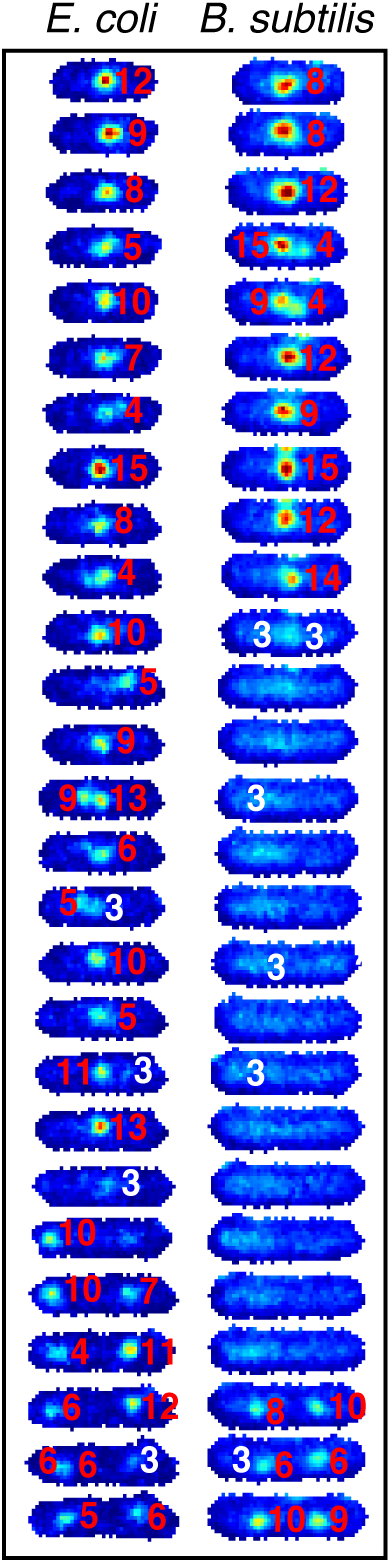
Automatic focus identification using scoring. Example scored foci in partial cell towers. Frame delay is five minutes. Scores are printed in red for foci scoring high enough to be included in analysis. Foci scoring 3 or lower (white) appeared randomly throughout the cell and were excluded.

**S3 Figure.**
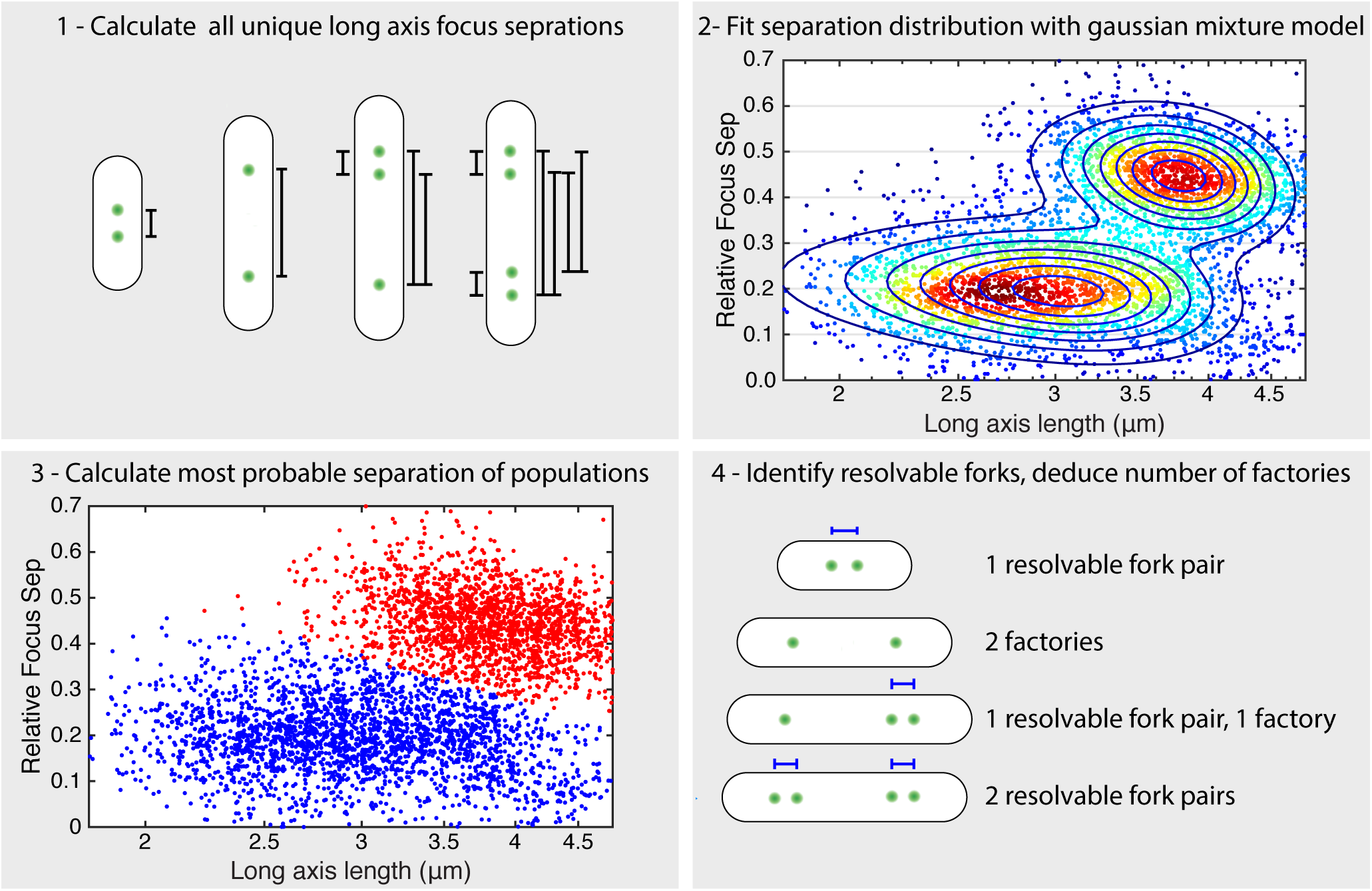
Process for distinguishing resolvable forks from pairs of factories in cells with multiple foci. 1) All possible long axis separations are calculated (black brackets). 2) Distribution of focus separation as a function of cell length (dots) is fit with a two Gaussian mixture model (dark blue contours) using maximum likelihood. We note that the fit to the lower population is shifted slightly due to the tail, however the populations separate visually correctly. 3) Using the Gaussian mixture model obtained in step 2, each focus pair is classified as a member of the high (red) or low (blue) separation population. 4) Focus pairs that are determined to be members of the low separation population (blue brackets) are classified as each representing an individual replisome. All other foci are inferred to be a colocalized replisome pair.

**S4 Figure.**
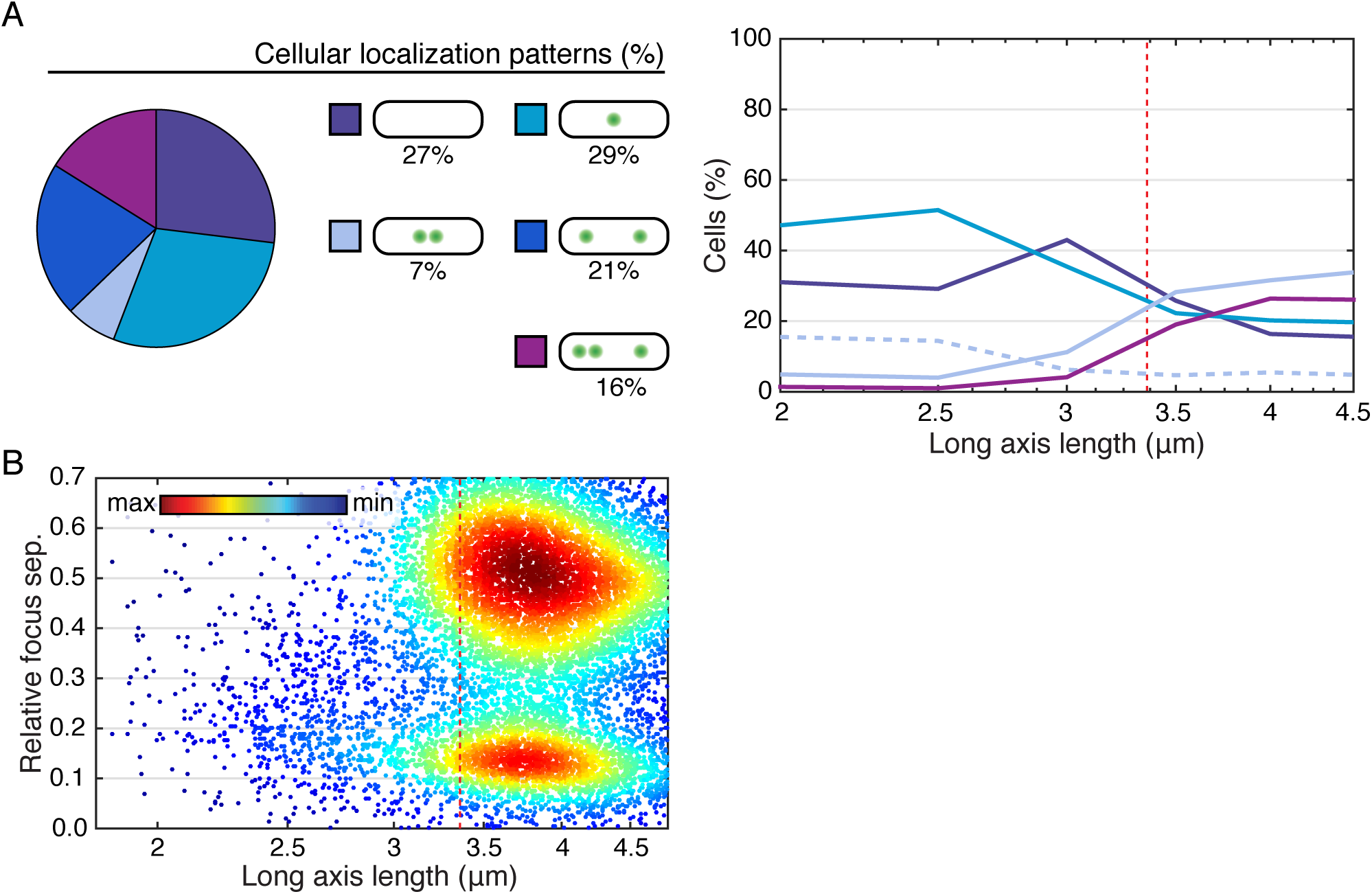
Supporting *B. subtilis* data. A) Relative frequencies of typical localization patterns overall (left) and separated by cell length (right). Dashed red line represents mean length at re-initiation B) Joint probability distribution for interfocus separation is used to automatically distinguish between cellular localization patterns. Dashed red line represents mean length at re-initiation

**S5 Figure.**
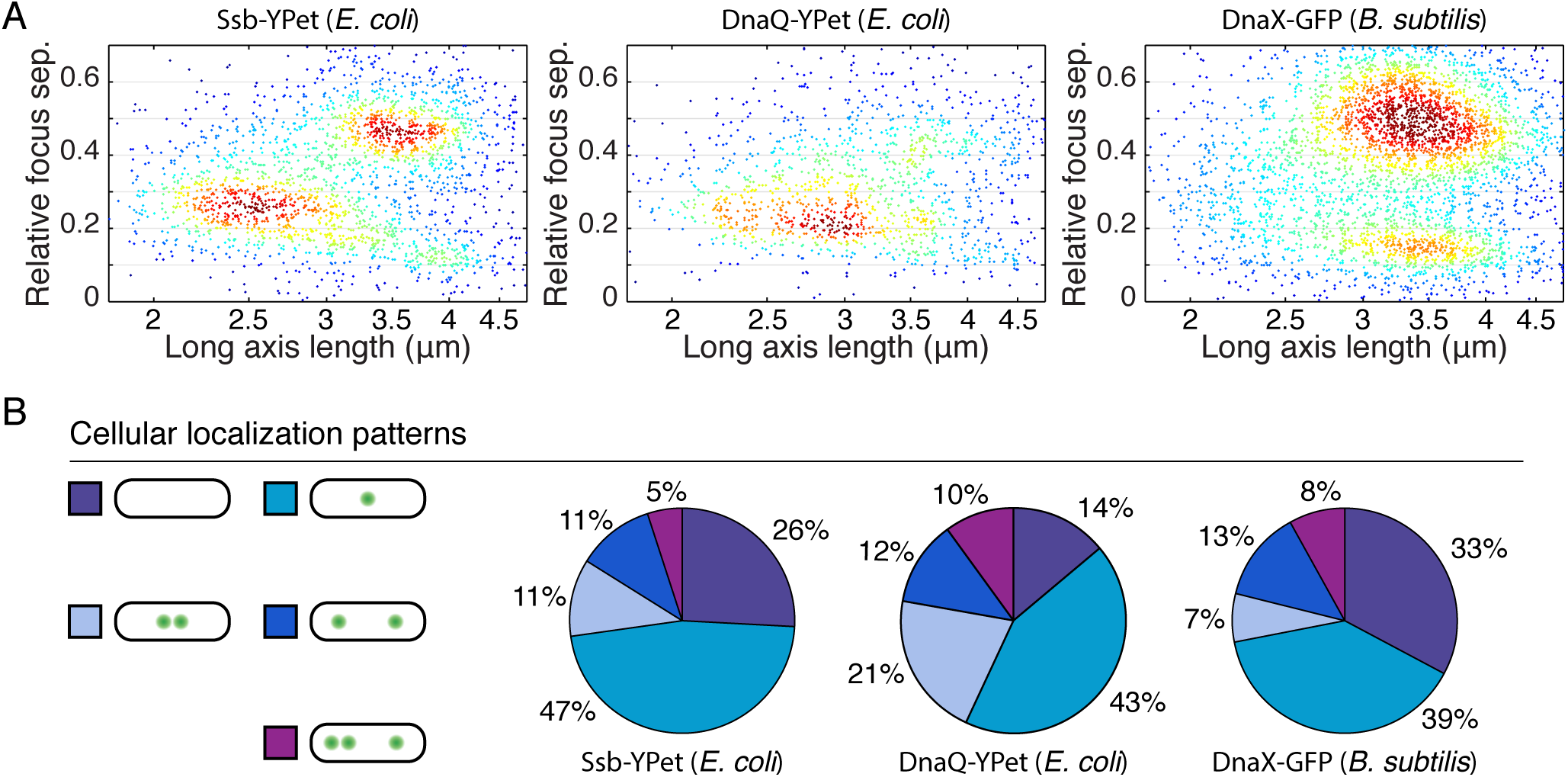
Snapshot analysis of several replisome proteins implies dynamics consistent with observations in time lapse imaging. A) Joint probability distributions for interfocus separation is used to automatically distinguish between cellular localization patterns in SSB-YPet (N = 6493 Cells), DnaQ-YPet (N = 2187 cells), and DnaX-GFP (N = 10573 cells). B) Relative frequencies of typical localization patterns.

## References

1. Lemon KP, Grossman AD (2000) Movement of replicating DNA through a stationary replisome. Molecular Cell 6(6):1321–1330.

2. Lemon KP, Grossman AD (1998) Localization of bacterial DNA polymerase: Evidence for a factory model of replication. Science 282(5393):1516–1519.

3. Molina F, Skarstad K (2004) Replication fork and SeqA focus distributions in *Escherichia coli* suggest a replication hyperstructure dependent on nucleotide metabolism. Molecular Microbiology 52(6):1597–1612.

4. Adachi S, Kohiyama M, Onogi T, Hiraga S (2005) Localization of replication forks in wild-type and mukB mutant cells of *Escherichia coli*. Molecular Genetics and Genomics 274(3):264–271.

5. Den Blaauwen T, Aarsman MEG, Wheeler LJ, Nanninga N (2006) Pre-replication assembly of *E. coli* replisome components. Molecular Microbiology 62(3):695–708.

6. Cass JA, Kuwada NJ, Traxler B, Wiggins PA (2016) *Escherichia coli* chromosomal loci segregate from midcell with universal dynamics. Biophysical Journal 110(12):2597–2609.

7. Bates D, Kleckner N (2005) Chromosome and replisome dynamics in *E. coli:* Loss of sister cohesion triggers global chromosome movement and mediates chromosome segregation. Cell 121(6):899–911.

8. Reyes-Lamothe R, Possoz C, Danilova O, Sherratt DJ (2008) Independent positioning and action of *Escherichia coli* replisomes in live cells. Cell 133(1):90–102.

9. Wallden M, Fange D, Lundius EG, Baltekin O, Elf J (2016) The synchronization of replication and division cycles in individual *E. coli* cells. Cell 166(3):729–739.

10. Goranov A, Breier A, Merrikh H, Grossman A (2009) YabA of *Bacillus subtilis* controls DnaA-mediated replication initiation but not the transcriptional response to replication stress. Molecular microbiology 74(2):454–466.

11. Migocki MD, Lewis PJ, Wake RG, Harry EJ (2004) The midcell replication factory in *Bacillus subtilis* is highly mobile: implications for coordinating chromosome replication with other cell cycle events. Molecular Microbiology 54(2):452–463.

12. Berkmen MB, Grossman AD (2006) Spatial and temporal organization of the *Bacillus subtilis* replication cycle. Molecular Microbiology 62(1):57–71.

13. Soufo CD1, Soufo HJ NGMSANPGP (2008) Cell-cycle-dependent spatial sequestration of the dnaa replication initiator protein in bacillus subtilis. Developmental Cell 95(1):935–941.

14. Kuwada NJ, Traxler B, Wiggins PA (2015) Genome-scale quantitative characterization of bacterial protein localization dynamics throughout the cell cycle. Molecular Microbiology 95(1):64–79.

15. Reyes-Lamothe R, Sherratt DJ, Leake MC (2010) Stoichiometry and architecture of active dna replication machinery in escherichia coli. Science 328(5977):498–501.

16. Kuwada NJ, Cheveralls KC, Traxler B, Wiggins PA (2013) Mapping the driving forces of chromosome structure and segregation in *Escherichia coli*. Nucleic Acids Research 41(15):7370–7377.

17. Stylianidou S, Brennan C, Nissen SB, Kuwada NJ, Wiggins PA (2016) Supersegger: robust image segmentation, analysis and lineage tracking of bacterial cells. Molecular Microbiology 102(4):690–700.

